# The Src family kinases Lck and Fyn promote transendothelial migration of T cells

**DOI:** 10.64898/2026.11.06.764914

**Authors:** Hope E Shearer, Nayeema Haque, Liz A Kurtz, Menelaos Symeonides, Nathan H Roy

## Abstract

The ability of T cells to cross endothelial barriers and migrate into and within peripheral tissue is crucial for adaptive immunity. Integrins such as LFA-1 play an essential role in this process by acting as adhesion molecules and initiating signaling events that drive the cytoskeletal changes needed for cell migration, however, many of the specific signaling events that promote T cell migration are still unknown. Here, we found that the Src family kinases Lck and Fyn are needed for efficient T cell migration. Primary T cells lacking Lck, Fyn, or both kinases exhibited defects in random migration on ICAM-1 coated surfaces for parameters such as speed and mean squared displacement. Analysis of signaling revealed that Lck was required for PI3K signaling downstream of LFA-1, while Fyn was not. Mechanistically, we found that the absence of Lck disrupted the normal interaction of the PI3K regulatory subunit p85 with tyrosine-phosphorylated binding partners, suggesting that Lck may regulate PI3K activity by controlling p85 protein-protein interactions. Importantly, both Lck and Fyn were required for efficient T cell transmigration across an inflamed endothelial barrier, a critical event for tissue infiltration. Lastly, a small-molecule Lck kinase inhibitor also decreased PI3K signaling and T cell transmigration, indicating the kinase activity of Lck is needed. Together, these data identify a new role for Lck and Fyn in promoting T cell transmigration.

## INTRODUCTION

The ability of T cells to migrate through vascular barriers is critical for a successful immune response to foreign threats, proper wound healing, and efficient tumor rejection. In response to inflammatory stimuli, local endothelial cells upregulate several factors, including adhesion molecules, chemokines, and cytokines. These inflammatory stimuli help recruit immune cells from the bloodstream and promote migration across the endothelial layer, a process called transendothelial migration (TEM)^1–4^. The integrin ligand ICAM-1 is highly upregulated on endothelial cells during inflammation and is critical for TEM of multiple immune cell types, including T cells, macrophages, neutrophils, and B cells ^5–10^. In T cells, the primary integrin that interacts with endothelial ICAM-1 is LFA-1. Interaction with endothelial ICAM-1 leads to firm adhesion to the endothelial wall and triggers signaling events that promote cytoskeletal and morphological changes that drive migration along and through the endothelial barrier ^8,11–15^. The importance of LFA-1 in normal T cell migration is exemplified by patients with a genetic deletion of LFA-1, causing severe immune cell migration defects and immunodeficiency ^16–18^.

The ligation of LFA-1 leads to the initiation of several downstream signaling events that drive T cell migration. These signaling events ultimately leads to cytoskeletal and morphological changes needed for migration, such as actin polymerization in the cell front to create protrusions ^11–13,19,20^. One important signaling pathway activated downstream of LFA-1 is the phosphoinositide 3-kinase (PI3K) pathway ^21–23^. PI3K is crucial for cell migration in many cell types and cellular contexts ^24^. The PI3K complex itself is composed of two subunits, a catalytic p110 subunit which converts phosphatidylinositol 4,5-bisphosphate (PIP_2_) into phosphatidylinositol 3,4,5-trisphosphate (PIP_3_), and a regulatory subunit which allosterically controls the activity of the p110 subunit. In general, local activity of PI3K and generation of PIP_3_ leads to the recruitment of cytoskeletal regulatory proteins (such as Rho GTPase family GEFs and GAPs), thus helping establish cell polarity and directional migration ^25,26^. Although LFA-1 ligation in T cells leads to PI3K activity ^21–23^, the specific signaling intermediates that link LFA-1 to PI3K remain largely unidentified.

The Src family kinases (SFKs) have been universally recognized as critical signaling intermediates downstream of integrins in numerous biological contexts, from development to cancer ^27,28^. The Src kinases are a small family of nonreceptor tyrosine kinases with nine classical members expressed in humans and only eight in mice: Src, Fyn, Yes, Lyn, Hck, Fgr, Blk, and Lck. SFKs generally localize to integrin adhesion sites where they signal through downstream effectors to influence pathways critical for cell function and survival ^29^. Immune cells vary in their expression of different SFKs (some of which are only expressed in immune cells, such as Hck, Fgr, and Lck), and their overlapping function makes it difficult to determine the roles of individual SFKs in migration. For instance, macrophages and neutrophils express Hck, Fgr, and Lyn, and deleting all three in macrophages disrupts adhesion and migration, while individual gene deletions have little to no impact ^30^. In neutrophil cell lines, some Lyn-specific roles in migration have been identified ^31^, and in primary neutrophils, manipulation of Hck and Fgr has yielded conflicting results ^32–34^. In T cells, the two predominantly expressed SFKs, Lck and Fyn, are heavily involved in proper T cell antigen receptor (TCR) signal transduction, which is critical for T cell development and activation ^35^. Because of their importance in TCR signaling and T cell activation and survival, it is difficult to study if these kinases play a role in primary T cell migration. For instance, mice lacking Lck have pronounced thymic atrophy and defective developmental signaling that reduces T cell populations ^36^, mice lacking Fyn have less severe but similar T cell survival impairments ^37^, and mice lacking both kinases develop no αβ T cells ^38^.

In this study, we found that the Src family kinases Lck and Fyn are crucial for efficient T cell migration. On ICAM-1-coated surfaces, primary T cells lacking Lck or Fyn displayed a lower mean squared displacement and migrated at slower speeds than WT T cells. In addition, our data show that T cells lacking Lck, but not Fyn, had severe defects in PI3K activation downstream of LFA-1 ligation. Mechanistically, Lck was needed for the normal association of PI3K p85 with tyrosine-phosphorylated partners. Inhibiting Lck pharmacologically also resulted in decreased PI3K activity, indicating that the kinase activity of Lck is needed to promote PI3K signaling. Finally, we asked if Lck and/or Fyn are needed for the ability of T cells to cross an inflamed endothelial barrier, a more complex directional migration situation with additional soluble and surface-bound contributing factors. Interestingly, knocking out either Lck or Fyn resulted in decreased transendothelial migration, suggesting Lck and Fyn are both needed for efficient T cell migration across inflamed endothelia. Together, these data identify a role for Src kinases in T cell migration.

## RESULTS

### T cells lacking the SFKs Lck and Fyn have defects in random migration on ICAM-1

Of the eight Src family kinases in the mouse genome, only two are predominantly expressed in αβ T cells, Lck and Fyn. To begin to test the role of Lck and Fyn in T cell migration, we first utilized a CRISPR/Cas9 system to generate primary CD4^+^ T cells lacking Lck, Fyn, or both ^39^. This CRISPR knockout (KO) protocol introduces gene editing after T cell activation, thus allowing for investigation of Lck and Fyn in migration independent of their known roles in T cell activation. Briefly, primary CD4^+^ T cells were isolated from secondary lymphoid tissue of Cas9-expressing mice, activated on anti-CD3 and anti-CD28 to induce proliferation, and then retrovirally transduced with either non-targeting (NT) or specific guide RNAs targeting Lck, Fyn, or both **(Fig. 1A)**. Protein levels of Lck and Fyn were significantly reduced in the individual KO conditions and in the dual KO condition (DKO) **(Fig. 1B-C)**. Importantly, loss of Lck, Fyn, or both did not impact the overall health and expansion of these T cells as measured by proliferation over time compared to the NT control **(Fig. 1D)**.

**Figure 1.**
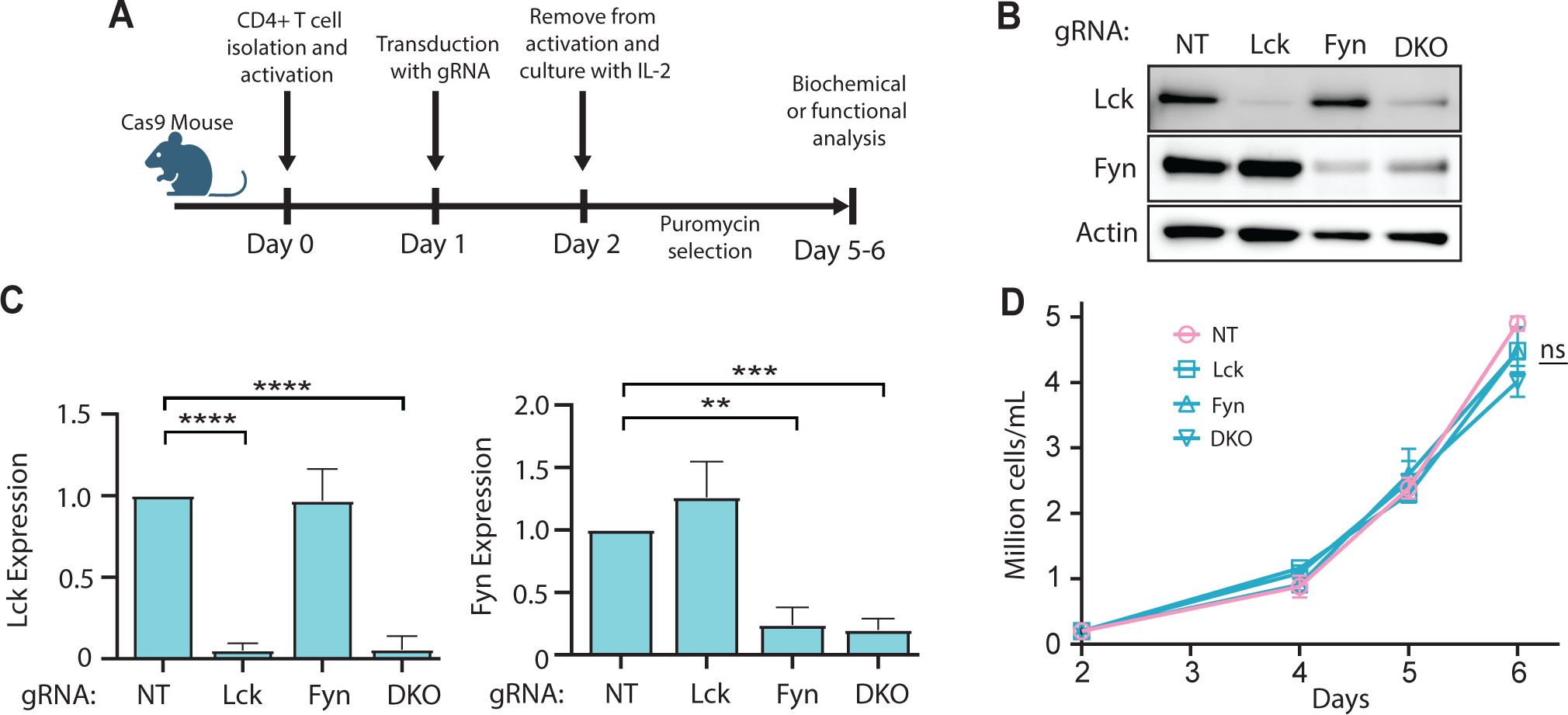
Generation of primary mouse CD4^+^ T cells lacking Lck and Fyn. **(A)** Workflow to generate CRISPR-mediated knockouts. **(B)** Western blots of lysates from non-targeting gRNA-transduced T cells (NT), Lck gRNA-transduced T cells (Lck), Fyn gRNA-transduced T cells (Fyn), and combined Lck and Fyn gRNA-transduced T cells (DKO). **(C)** Quantification of Lck and Fyn band intensities normalized to actin in each condition of n=3 independent experiments. **(D)** Growth curve comparing NT, Lck, Fyn, and DKO cell numbers in culture on days 2-6 after activation from n=3 independent experiments. *p<0.05, **p<0.01, ***p<0.001, ****p<0.0001 (ordinary one-way ANOVA with multiple comparisons post-test). Error bars represent SD.

To determine if Lck and/or Fyn contribute to T cell migration, we allowed NT, Lck KO, Fyn KO, or DKO T cells to randomly migrate on ICAM-1-coated chambers and imaged them using time-lapse microscopy. The cells were imaged at low magnification for 10 min and then tracked using FiJi ^40^. The percentage of migratory cells (defined as cells that moved more than two cell lengths during the imaging period) did not vary significantly between groups, indicating the KO cells were capable of successful adherence and motility **(Fig. 2A)**. We then individually tracked migratory cells and exported x-y coordinates for each time point into the open-source analysis software Migrate3D ^41^.

**Figure 2.**
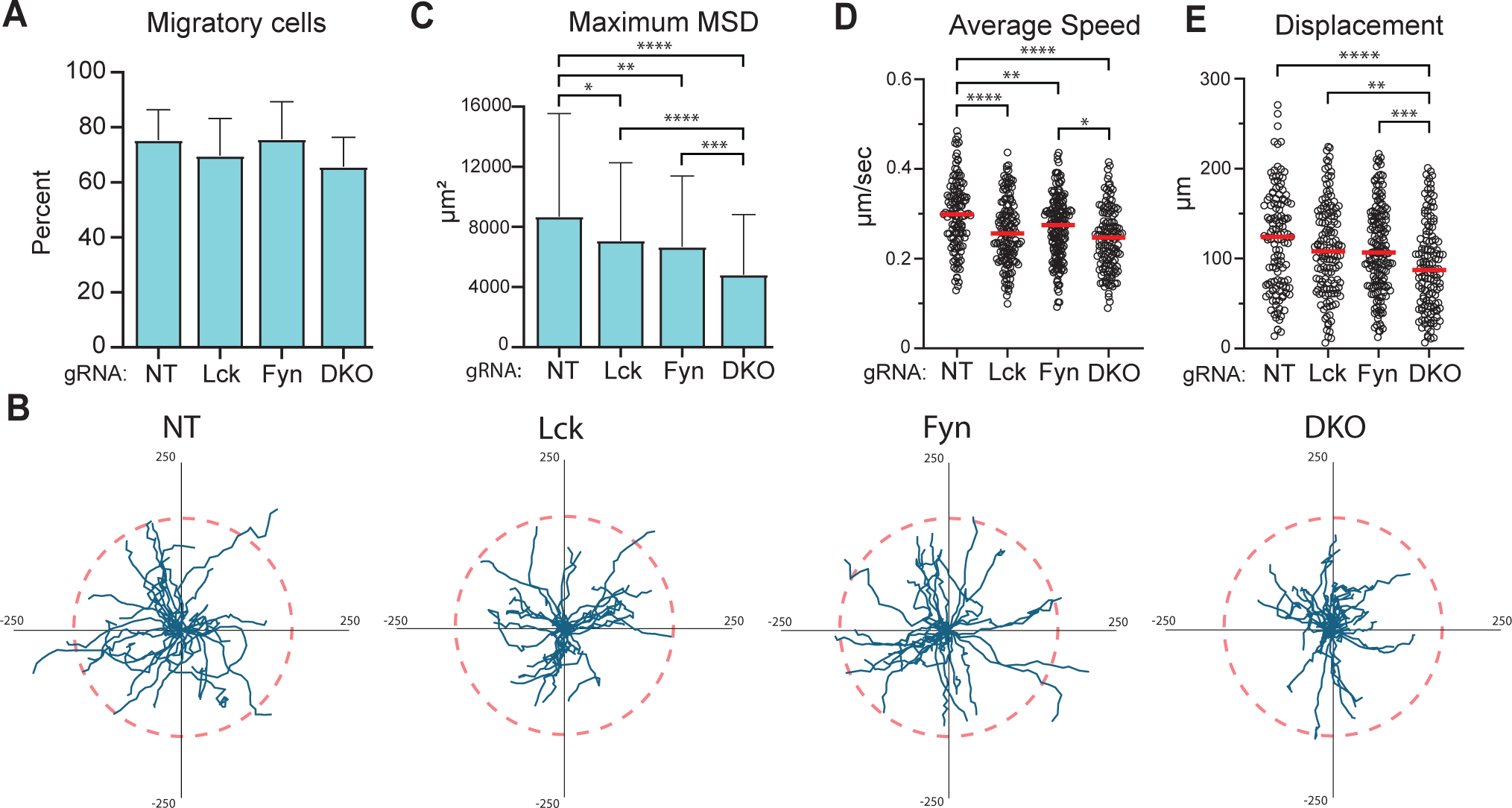
Lck, Fyn, and double knockout T cells exhibit defects in random migration on ICAM-1. **(A)** T cells were allowed to migrate on ICAM-1 coated surfaces and imaged for 10 min. T cells were then tracked and the percentage of migratory cells in each condition was calculated (defined by the fraction of cells that move more than 2 cell lengths during the imaging period). Pooled from n=3 experiments (cell totals: 242 NT, 233 Lck KO, 284 Fyn KO, and 301 DKO). **(B)** Spider plots of 30 cell tracks from a representative experiment. All tracks were quantified for **(C)** maximum mean squared displacement (MSD), **(D)** average speed, and **(E)** displacement. Data pooled from n=3 independent experiments with 127 NT cells, 142 Lck KO cells, 170 Fyn KO cells, and 141 DKO cells. *p<0.05, **p<0.01, ***p<0.001, ****p<0.0001 (ordinary one-way ANOVA with multiple comparisons post-test). Error bars represent SD.

Spider plots of a representative experiment show a slightly tighter migratory spread of the Lck KO and DKO T cells compared to the NT control (**Fig. 2B**). For quantitative analysis, we used Migrate3D to calculate several migration parameters, including maximum mean squared displacement (MSD), speed, and displacement (**Fig. 2C-E**). T cells lacking Lck, Fyn, or both had significantly lower maximum MSD and speed compared to the non-targeting control (**Fig. 2C-D**). Interestingly, DKO T cells were even more defective than the single KO counterparts in both MSD and overall displacement **(Fig. 2C, E)**. These data indicate Lck and Fyn have an important role in maintaining proper T cell migration on ICAM-1.

### T cells lacking SFKs exhibit defects in crossing inflamed endothelia

T cell transmigration across inflamed endothelia is highly dependent on integrin-ligand interactions ^2^. However, many other factors can influence TEM, including other adhesion molecules and chemokines that are expressed during inflammation. To determine the importance of SFKs in this more relevant, complex migration process, we utilized an *in vitro* microscopy-based assay to measure the ability of T cells to cross an inflamed endothelium. 3B-11 mouse endothelial cells were grown to confluency and treated with TNFα to mimic an inflammatory state. As shown in **Figure 3A**, treatment with TNFα drastically increased surface ICAM-1 expression on the endothelial layer. To measure how TNFα treatment impacts T cell TEM, we allowed WT T cells to settle on untreated or TNFα-treated endothelial cells and imaged for 1 h. The cells captured within the 1 h movies were first classified as migratory or non-migratory (as in **Fig. 2A**), and then the migratory cells were scored on whether they crossed the endothelium or remained on the apical side **(Fig. 3B-C, Supp Movie 1)**. Quantification of several hundred cells over multiple experiments shows that about 60% of T cells transmigrate under these conditions (**Fig. 3B**). To determine if the SFKs play a role in TEM, we allowed Lck, Fyn, and double KO T cells to migrate on TNFα-treated endothelial cells and measured transmigration as above. T cells lacking Lck, Fyn, or both kinases had significant defects in TEM (**Fig. 3D**). These data indicate that SFKs play a critical role in T cell migration across inflamed endothelium.

**Figure 3.**
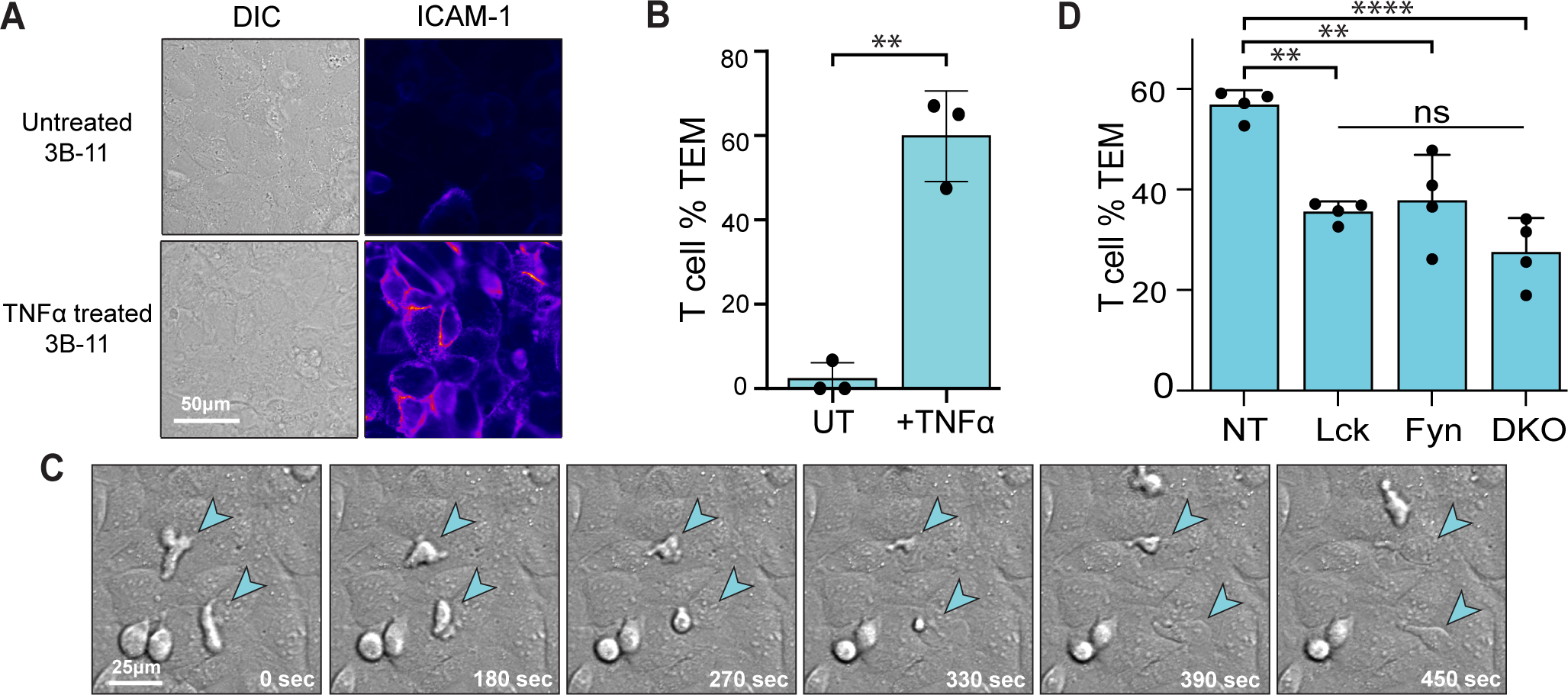
SFKs are needed for transmigration across inflamed endothelia. **(A)** 3B-11 endothelial cells were left untreated or treated with TNFα (50 ng/mL) for 24 h, then surface stained with anti-ICAM-1. **(B)** T cells were allowed to migrate on endothelia, imaged every 30 seconds for 1 h, and TEM was quantified. Data pooled from n=3 independent experiments with 163 T cells on untreated 3B-11s and 227 T cells on TNFα-treated 3B-11s. **(C)** Representative TEM events. **(D)** Quantification of TEM of NT, Lck KO, Fyn KO, or DKO T cells on TNFα-treated endothelia. n=4 independent experiments with 462 NT, 505 Lck KOs, 672 Fyn KOs, and 721 DKOs. *p<0.05, **p<0.01, ***p<0.001, ****p<0.0001 (ordinary one-way ANOVA with multiple comparisons post-test). Error bars represent SD.

### T cells lacking Lck have blunted PI3K activation

Several signaling pathways are important for proper T cell migration. We and others have found primary mouse T cells robustly activate PI3K while migrating on the LFA-1 ligand ICAM-1, as measured by phosphorylation of AKT ^20,23^. To test if SFKs are involved in PI3K signaling in response to migration on integrin ligands, NT or KO T cells were allowed to settle on ICAM-1-coated surfaces for 20 min, lysed, and immunoblotted for phosphorylated AKT at T308 for a direct readout of PI3K activation and S473 as a readout of both PI3K and mTORC2 activity **(Fig. 4A-B)**. As expected, NT T cells showed robust phosphorylation of AKT at both sites after LFA-1 ligation, however, Lck KO and DKO T cells showed significantly less phospho-AKT at both sites **(Fig. 4A-B)**. Interestingly, the untreated Fyn KO T cells had higher baseline phospho-AKT at site T308 than NT, Lck KO, and DKO T cells, however after ICAM-1 stimulation, the levels of T308 were not significantly different compared to the NT control.

**Figure 4.**
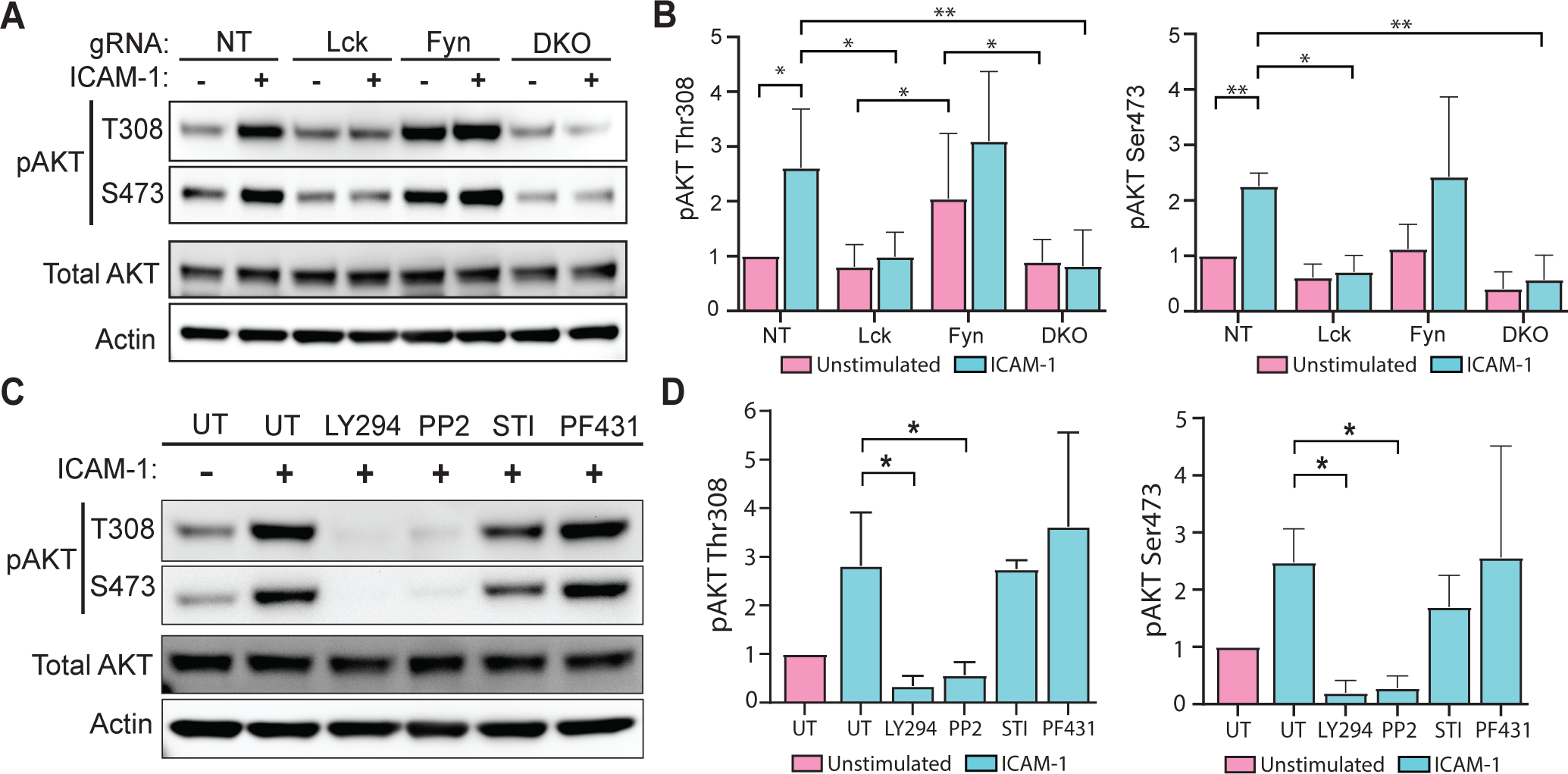
Lck is needed for PI3K activation in migrating T cells. **(A)** T cells transduced with the indicated gRNA were allowed to migrate on ICAM-1 for 20 min, or left unstimulated as a control, lysed, and immunoblotted with the indicated antibodies. **(B)** Quantification of the pAKT T308 and S473 signal normalized to unstimulated NT control. Minimum of n=4 independent experiments per condition. **(C)** Western blot of lysates from WT T cells allowed to migrate on ICAM-1 for 20 min either untreated (UT) or treated with the indicated kinase inhibitor (PI3K inhibitor LY294002 (20 μM), Src family kinase inhibitor PP2 (10 μM), Ablson family kinase inhibitor STI/Imatinib (10 μM), or focal adhesion kinase family inhibitor PF431396 (5 μM)) and immunoblotted for pAKT T308 and S473. **(D)** Quantification of pAKT T308 and S473 signal normalized to unstimulated UT control from n=3 independent experiments. *p<0.05, **p<0.01 (ordinary one-way ANOVA with multiple comparisons post-test. Error bars represent SD.

To determine if other kinase families known to be important for integrin signaling also control PI3K activity in T cells, we utilized a panel of tyrosine kinase inhibitors and measured pAKT. Our inhibitor panel included PP2 (broad-spectrum Src family), PF431 (FAK family), STI-371 (Abelson family), and a PI3K inhibitor control, LY294002. T cells were allowed to settle on ICAM-1-coated surfaces in the presence or absence of the indicated inhibitor for 20 min (there was no pre-treatment with inhibitors), lysed and immunoblotted for pAKT at both T308 and S473, as done above **(Fig. 4C-D)**. As expected, the positive control LY294002 blocked phosphorylation at both these sites. Of our experimental inhibitors, only the Src kinase inhibitor PP2 caused blunted phosphorylation of AKT at T308 and S473, while the Abl family inhibitor STI-371 and the FAK family inhibitor PF-431396 had no effect **(Fig. 4C-D)**. Together, these data indicate that Lck is needed for AKT phosphorylation, and thus PI3K activity, in T cells migrating on ICAM-1.

### Loss of Lck disrupts the integration of p85 with tyrosine-phosphorylated partners

The catalytic subunit of PI3K, p110, is controlled by the regulatory subunit, p85. When p85 binds to pTyr residues, it triggers a structural change that releases the inhibition of the catalytic subunit p110, resulting in enzymatic activity and the production of PIP_3_ ^42^. Previous work has identified that p85 interacts with several prominent tyrosine-phosphorylated proteins when T cells migrate on ICAM-1 ^20^. Therefore, we asked if Lck is needed for the normal association of p85 with phosphorylated partner proteins during migration. First, we used a broad anti-pTyr antibody to probe the total pTyr changes in NT and KO T cells migrating on ICAM-1 or left unstimulated as a control **(Fig. 5A)**. Lck KO and DKO T cells displayed clear losses in pTyr signal at multiple molecular weights, while Fyn KO T cells remained unchanged in pTyr signal **(Fig. 5A)**. Quantification of the densest region around 120 kDa showed that Lck and DKO T cells had an almost complete loss of pTyr signal **(Fig. 5B)**. Previous work shows that p85 interacts with a tyrosine-phosphorylated protein of this size that was identified by mass spectrometry as c-Cbl ^20^. To determine if the SFKs control normal p85 binding to tyrosine-phosphorylated proteins such as c-Cbl, we immunoprecipitated p85 from NT and KO T cells after migrating on ICAM-1 (or left unstimulated as a control) and immunoblotted for pTyr **(Fig. 5C)**. A tyrosine-phosphorylated protein co-immunoprecipitated with p85 at ∼120 kDa in NT T cells only after migrating on ICAM-1. However, in Lck KO and DKO T cells, p85 failed to co-immunoprecipitate this pTyr species **(Fig. 5C)**. Conditions in which p85 co-immunoprecipitated a pTyr protein at ∼120 kDa also showed enrichment of c-Cbl by immunoblotting, suggesting that the pTyr species could correspond to c-Cbl. These data indicate that Lck is needed for the normal association of p85 with tyrosine-phosphorylated binding partners.

**Figure 5.**
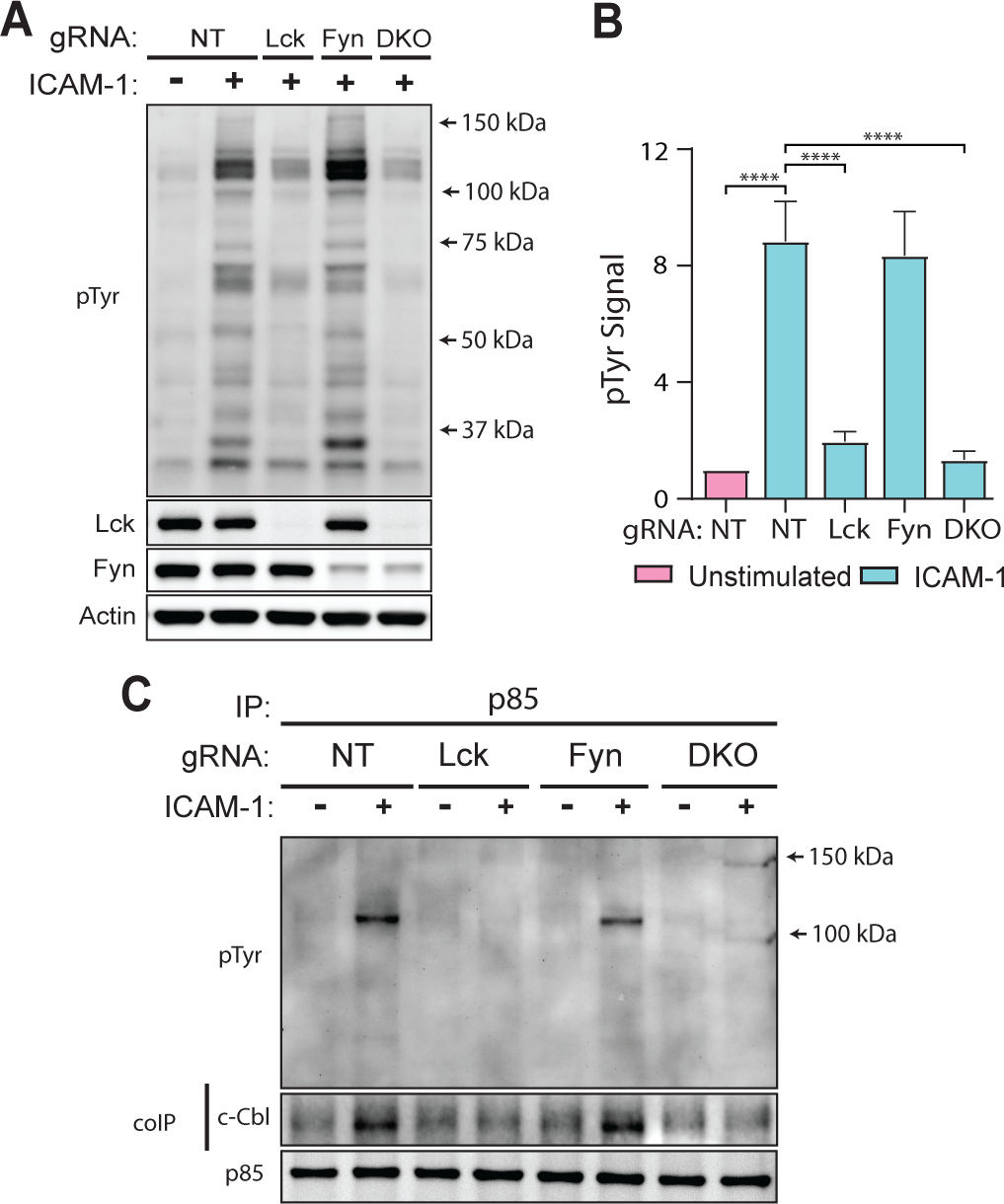
The absence of Lck alters tyrosine-phosphorylated binding partners of p85. **(A)** NT and KO T cells allowed to migrate on ICAM-1 for 20 min were lysed and immunoblotted with the indicated antibodies. **(B)** Quantification of pTyr signal normalized to the NT control at the 120kDa region for each condition. n=3 independent experiments. **(C)** NT and KO T cells were allowed to migrate on ICAM-1 for 20 min, or left unstimulated as a control, then lysed and immunoprecipitated for p85. Samples were immunoblotted for pTyr, p85, and c-Cbl. Representative of n=3. ****p<0.0001 (ordinary one-way ANOVA with multiple comparisons post-test). Error bars represent SD.

### Lck catalytic activity is needed to promote PI3K signaling and TEM

Lck is a multi-domain protein that acts as a tyrosine kinase and can also function as an adaptor in some situations ^43^. Therefore, we asked if the kinase activity of Lck was needed to promote PI3K activity and transmigration in migrating T cells. To do this, WT T cells were allowed to migrate on ICAM-1 coated surfaces in the presence or absence of the Lck inhibitor RK24466 for 20 min (no pre-treatment), lysed and immunoblotted for p-AKT. Migrating T cells treated with a high concentration (2.5 μM) or a low concentration (40 nM) of RK24466 had significantly less phosphorylation of AKT at T308 and S473 **(Fig. 6A-B)**. The broad-spectrum SFK inhibitor PP2 was utilized as a positive control. To determine if Lck kinase activity was needed for crossing inflamed endothelia, we repeated the TEM experiments in the presence or absence of RK24466. Addition of RK24466 for the 1 h imaging period significantly reduced TEM **(Fig. 6C)**. These data suggest that the kinase activity of Lck is needed to promote PI3K signaling and TEM of T cells.

**Figure 6.**
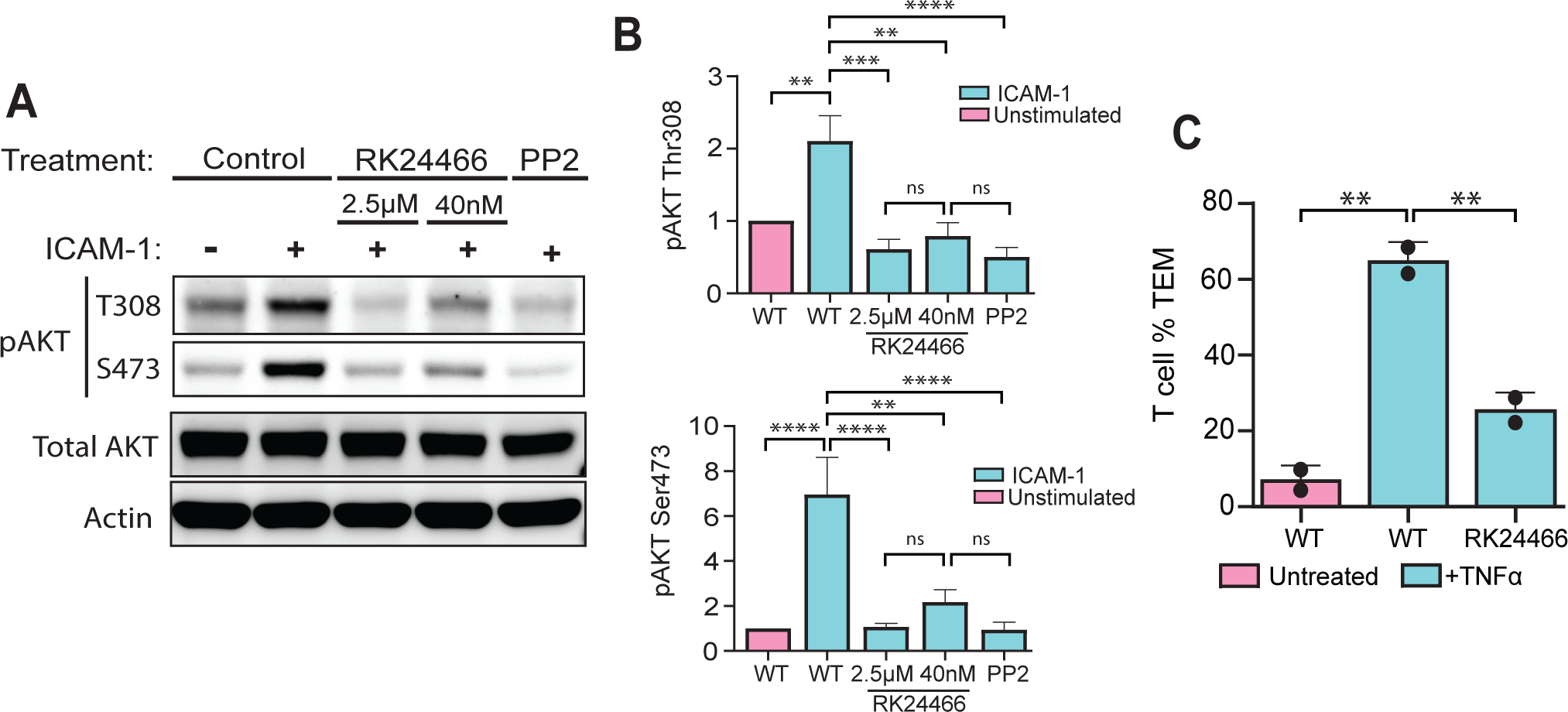
The Lck inhibitor RK24466 disrupts AKT phosphorylation and transmigration. **(A)** WT T cells were allowed to migrate on ICAM-1 for 20 min while simultaneously treated with the Lck inhibitor RK24466 at either 2.5 μM, 40 nM, or left untreated as a control. The broad SFK inhibitor PP2 (10 μM) was also included as a positive control. Cells were lysed and immunoblotted with the indicated antibodies. **(B)** Quantification of AKT phosphorylation at sites S473 and T308. n=3 independent experiments. **(C)** Quantification of TEM in untreated or RK24466 treated (40 nM), on either untreated or TNFα-treated 3B-11 cells. n=2 with 64 T cells on untreated 3B-11s, 103 T cells on TNFα-treated 3B-11s, and 156 RK24466-treated T cells on TNFα-treated 3B-11s. *p<0.05, **p<0.01, ***p<0.001, ****p<0.0001 (ordinary one-way ANOVA with multiple comparisons post-test). Error bars represent SD.

## DISCUSSION

Past studies have uncovered distinct roles for kinases such as FAK1, PYK2, EphA1, and WNK1 in LFA-1 signaling and T cell adhesion and migration ^19,44–46^. In addition, work in T cell lines has identified several other kinases that are active downstream of LFA-1 and contribute to T cell migration, including ZAP-70 and Casein Kinase 2 ^47–49^. In this study, we reveal that the SFKs Lck and Fyn both contribute to T cell migration. Primary T cells lacking either kinase migrated more slowly and had decreased mean squared displacement on ICAM-1 coated surfaces. Importantly, T cells lacking either Lck or Fyn also displayed significant defects in crossing an inflamed endothelial monolayer.

Evaluation of signaling revealed that Lck, but not Fyn, was crucial for PI3K signaling in migrating T cells. Mechanistically, we found that Lck is needed for the normal association of the PI3K subunit p85 with tyrosine phosphorylated proteins, suggesting that Lck may control the activity of PI3K by regulating its binding partners. This function is likely directly due to the kinase activity of Lck, as a targeted kinase inhibitor also inhibited PI3K signaling and T cell TEM. Together, these results define a new role for the SFKs in T cell migration.

Most work on Lck and Fyn in primary T cells has focused on their central roles in T cell activation and expansion ^35^. The role of Lck and Fyn in migration is less clear, for instance, some studies in T cell lines have documented that Lck is needed for chemotaxis, while others indicate that Lck is dispensable and that Fyn is needed ^50–56^. Because T cell lines contain numerous mutations and abnormalities that affect normal cell signaling and migration, we used an approach to disrupt Lck and Fyn in activated primary T cells to investigate their role in migration *ex vivo*. Our data indicate that both Lck and Fyn contribute to T cell migration on the integrin ligand ICAM-1 **(Fig. 2)**. T cells lacking Lck, Fyn, or both displayed decreases in mean squared displacement, speed, and directionality, all important measures of efficient T cell migration. While SFKs are well known to localize to focal adhesion complexes and regulate adhesion and migration in non-hematopoietic cells ^27–29^, adhesion complexes have been difficult to study in T cells. However, a recent report indicates that primary T cells can form small, transient adhesion complexes under some conditions ^57^. In line with their role in other systems, Lck and/or Fyn may regulate T cell migration by physically localizing to these small adhesion complexes to influence downstream signaling.

Our study identifies Lck and Fyn as necessary for efficient transmigration of T cells across inflamed endothelia. In other immune cells such as monocytes and neutrophils, the SFKs Hck, Fgr, and Lyn are needed for efficient TEM and likely have overlapping functions ^58,59^. It has been less clear if SFKs play a role in T cell transmigration, for instance, studies using the broad SFK inhibitor PP2 have shown mixed results depending on the experiment ^8,60^. One study indicates that SFKs are important for transcellular diapedesis, but not paracellular diapedesis of T cells ^61^. Here, our more pointed genetic approach showed that KO of Lck, Fyn, or both, in primary T cells result in significant defects in TEM. Interestingly, DKO T cells had similar defects in TEM as each single Lck or Fyn KO, suggesting that 1) Lck and Fyn cannot compensate for the function of the other in regulating TEM and 2), Lck and Fyn may function in the same pathway or exert their effects on a similar aspect of migration. These results are somewhat at odds with the data indicating that loss of Lck affects PI3K signaling, but loss of Fyn does not (**Fig. 4**). While we attribute the TEM defects in Lck KO T cells (at least partially) to the loss of PI3K activity, it is still unclear the potential mechanisms in which Fyn may promote TEM. It is worth noting that untreated Fyn KO T cells do display increased PI3K signaling at baseline, suggesting Fyn may negatively regulate PI3K activity (opposing Lck), however, when migrating on ICAM-1 there was no difference from NT control T cells. It is possible our immunoblot assay is not sensitive enough to determine a true difference, as one study did reveal that Fyn can negatively regulate Lck activity in T cells ^62^. Nevertheless, if these proteins do work in the same pathway, as our TEM data suggest, Fyn could instead function downstream of PI3K activity, for instance regulating some of the cytoskeletal processes triggered by PI3K that are required for migration. Future experiments will explore these potential mechanisms.

One major mechanism of PI3K regulation is the allosteric inhibition of enzymatic activity by the regulatory subunit p85. Binding of the p85 regulatory subunit to tyrosine-phosphorylated proteins relieves this inhibition and results in structural changes and p110 catalytic activity ^42^. Previous work has found that p85 interacts with a highly phosphorylated protein around 120 kDa, which was identified by mass spectrometry as the protein c-Cbl (and smaller amounts of the Cbl family member Cbl-b) ^20^. Here, we found that Lck was needed for the normal interaction of p85 with phosphorylated partners during T cell migration (**Fig. 5**). Several studies in cell lines have suggested that Lck is involved in PI3K signaling downstream of surface receptors such as the TCR ^63–66^. The potential involvement of Lck in migration has been difficult to study in primary T cells due to the central role of Lck in T cell activation. Our data suggest a mechanism in which Lck phosphorylates a binding partner(s) for p85 (likely c-Cbl or Cbl-b), causing relief of PI3K autoinhibition and catalytic activity.

One study proposed a similar mechanism involving Lck phosphorylation of the receptor Tim-1, which then recruits p85 causing localized PI3K activity ^67^. It is possible that the identified p85 binding partner in our study also serves as an adaptor to help localize active PI3K to the cell front or to where protrusions are needed. The substrates of Lck during T cell migration, and their effects on PI3K activity and migration will be the focus of future work.

In conclusion, we have identified that the Src family kinases Lck and Fyn are needed for efficient T cell transmigration across inflamed endothelia. Our data indicate that these kinases have unique roles in promoting transmigration, and also strengthen the idea that Lck and Fyn have clear functions beyond T cell activation and development in primary T cells.

## ACKNOWLEDGMENTS

We thank the SUNY Upstate Medical University Department of Laboratory Animal Resources for providing care to our mouse colony. We also thank the members of the Roy laboratory, Liz Kurtz, Nayeema Haque, Alex Fischette, and Vamsi Adabala, for the discussions, moral support, and critical reading of the manuscript.

## MATERIALS AND METHODS

### Antibodies and reagents

Anti-MHCII clone M5/114, anti-CD8 clone 2.43, anti-CD3 clone 2C-11, anti-CD54 (ICAM-1) clone YN1/1.7.4 and anti-CD28 clone PV-1 were obtained from BioXCell. Western blot and immunofluorescence antibodies were: from Cell Signaling, anti-actin clone 13E5 (cat. # 4970S), anti-Lck clone D88 (cat. # 2984S), anti-Fyn (cat. # 4023S), anti-pAKT(S743) clone D9E (cat. # 4060S), anti-pAKT(T308) clone D25E6 (cat. # 13038S), anti-AKT (pan) clone 40D4 (cat. # 2920S), anti-p85 (cat. # 4292S), and anti-c-Cbl (cat. # 2747S). Anti-pTyrosine clone PY20 (product # 05-947) was purchased from Sigma-Aldrich. Anti-Lck clone A17013D (cat. # 601002) was purchased from BioLegend. We obtained secondary antibodies conjugated with appropriate fluorophores or horseradish peroxidase from Jackson ImmunoResearch or Life Technologies. Recombinant mouse ICAM-1 was purchased from R&D Systems. Small molecule inhibitors PP2 (cat # 13198), LY294002 (cat. # 70920), STI-571 (cat # 13139), and RK24466 (cat # 15135) were purchased from Cayman Chemicals. The recombinant mouse TNFα was purchased from R&D biotechne (cat # 410-MT-025/CF). The Cytiva Amersham ECL western blotting detection reagent was purchased through Thermo Fisher Scientific (cat # 45-002-401).

### Mice

All mouse husbandry was done in the Upstate Medical University mouse facility in accordance with the Institutional Animal Care and Use Committee guidelines. WT B6 (C57BL/6J, strain #000664) and Cas9 (B6J.129(Cg)-Igs2tm1.1(CAG−cas9*)Mmw/J, strain #028239) mice were purchased from The Jackson Laboratory. All mice were 8–16 weeks of age when cells were harvested, and both male and female mice were used equally as sources of T cells for *in vitro* assays.

### Cell Culture

For in vitro assays, primary mouse CD4^+^ T cells were isolated from spleens and lymph nodes from 8–16-week-old mice. Briefly, tissues were homogenized and filtered before incubation with ACK lysis buffer (77 mM NH_4_Cl, 5 mM KHCO_3_, and 55 μM EDTA, pH 7.2–7.4) for 1 min to remove red blood cells (RBCs). After RBC lysis, cells were washed and incubated with anti-CD8 (1 μg/mL) and anti-MHCII (1 μg/mL) for 20 min at 4°C. After initial incubation, cells were washed and mixed with BioMag goat anti-rat IgG beads (Qiagen cat. # 310107) and incubated for an additional 15 min at 4°C before samples were placed onto a magnetic rack for three rounds of magnetic separation. Isolated CD4^+^ T cells were then counted and immediately plated on a 24-well plate coated with 5 μm/mL anti-CD3 and 2 μm/mL anti-CD28 at 10^6^ cells per well. Activation was done in complete T cell medium composed of Corning DMEM supplemented with 5% FBS, 1% each of penicillin-streptomycin, non-essential amino acids, and Glutamax (all purchased from Gibco), plus 2 μL 2-mercaptoethanol (obtained from Sigma-Aldrich). After 48 h of activation, T cells were removed from activation and mixed at a 1:1 volume ratio with complete T cell medium containing recombinant IL-2 (obtained from Sigma-Aldrich) to a final concentration of 20 U/mL IL-2. T cells were used on days 5–6 post-harvest for all experiments.

293T cells (for retroviral production) were originally obtained from ATCC, maintained in DMEM supplemented with 10% FBS, 1% nonessential amino acids (Gibco), and Glutamax (Gibco), and regularly screened for mycoplasma.

3B-11 endothelial cells (for TEM assays) were obtained from ATCC and were maintained in DMEM supplemented with 10% FBS, Glutamax (Gibco), and 150 mg/L of sodium bicarbonate.

### Retroviral production

Specific gRNAs to Lck and Fyn were cloned into the retroviral transfer vector MRIG ^39^. This vector was previously modified to express a puromycin resistance gene in place of the original GFP for downstream selection ^68^. The gRNA sequences used in this study were as follows: non-targeting (NT) 5-GCGAGGTATTCGGCTCCGCG-3; Lck 5’-TGAGCCGTAAGGACGCCGAG-3’; and Fyn 5’-AGCCCGCTCCTTGACAACCG-3’. To produce retroviral particles containing guide RNAs targeting Lck and Fyn, 293T cells were grown to 50% confluency, then co-transfected with 4.5 μg MRIG-puro along with 3.5 μg pCL-eco packaging vector (Addgene #12371). Transfection was done using calcium phosphate purchased from Invitrogen (cat. # K278001), following the manufacturer’s protocol.

Transfection mixes remained on 293T cells overnight before being replaced with fresh medium, after which cells were incubated for an additional 24–30 h. Retrovirus-containing supernatants were collected and centrifuged for 10 min at 1000 g to remove cellular debris, and polybrene (Millipore Sigma) was added at a final concentration of 8 μg/mL. Retroviral supernatants were then immediately applied to T cells for transduction.

### T cell transduction

CD4^+^ T cells that were isolated from Cas9-expressing mice were activated on 24-well plates coated with 5 μg/mL anti-CD3 and 2 μg/mL anti-CD28 at 10^6^ cells per well in complete T cell medium. After 24 h of activation, the T cell medium was collected and reserved for post-transduction, and 800 μL of viral supernatant was gently added per well before the plates were incubated at 37°C for 10 min. Following incubation, plates were centrifuged at 1100 g for 2 h at 35°C. Directly following spinoculation, plates were incubated at 37°C for 10 min; viral supernatants were removed and replaced with 1.5 mL of conditioned medium mixed with fresh complete T cell medium at a 2:1 volume ratio. T cells were cultured for an additional 24 h before being removed from activation and mixed at a 1:1 volume ratio with complete T cell medium containing recombinant 40 U/mL IL-2 to create a final concentration of 20 U/mL. After 2 h incubation in IL-2 medium at 37°C, medium containing puromycin was added so that the final concentration was 4 μg/mL. Transduced cells were used for assays on day 5 and 6 post-isolation.

### Biochemical analysis of T cell signaling in response to surface-bound ICAM-1

60 mm tissue culture dishes were coated with 2 μg/mL ICAM-1 overnight at 4°C. Activated CD4^+^ T cells were serum-starved in FBS-free DMEM for 2 h at 37°C then washed and resuspended in warm HBSS +/+ (Gibco) supplemented with 2 mg/mL D-glucose (Millipore Sigma). Cells were incubated for 10 min at 37°C before 10 million cells were added to the 60 mm dishes for 20 min or left unstimulated as a control. ICAM-1-stimulated lysates were collected by placing dishes on ice and adding lysis buffer (1% Triton X-100, 150 mM NaCl, 50 mM Tris-HCl pH 7.5, 5 mM NaF, 1 mM sodium orthovanadate, and Roche EDTA-free protease inhibitor). Unstimulated control cells were lysed in suspension by placing them on ice and adding cold 2x lysis buffer at a 1:1 volume ratio. Lysates were kept on ice and vortexed periodically for 10 min, followed by centrifugation at 13,000 g for 10 min at 4°C. Clarified whole-cell lysate was mixed with 4x sample buffer (Invitrogen) with added DTT (Roche, 50 mM final concentration) and heated for 8 min at 95°C before separation with SDS-PAGE.

### Western blotting

Proteins were separated by SDS-PAGE using the Invitrogen Novex Mini-cell system with NuPAGE 4-12% BisTris gradient gels. Proteins were transferred to a nitrocellulose membrane (0.22 μm, Cytex) and blocked for 1 h at room temperature with Li-Cor blocking buffer mixed 1:1 (v/v) with TBS. Primary antibodies were mixed in TBST (50 mM Tris-HCl pH 7.4, 150 mM NaCl, and 0.1% Tween-20) with 2% BSA and added to membranes overnight at 4°C. Primary antibody dilutions were as follows: anti-Actin (1:10,000), anti-Lck (1:1,000), anti-Fyn (1:1,000), anti-pAKT S473 (1:1,000), anti-pAKT T308 (1:500), anti-AKT (1:1,000), anti-p85 (1:1,000), anti-c-Cbl (1:1,000), anti-pTyr (1:500). Membranes were washed three times with TBST for 5 min each wash before the addition of fluorophore-conjugated secondary antibodies diluted in TBST with 2% BSA and incubated at room temperature for 1 h.

Membranes were then washed three times with TBST for 5 min each wash, then transferred into PBS for imaging on the ChemiDoc Imaging System (Bio-Rad). Band intensities were quantified using ImageJ software. If horseradish peroxidase (HRP) was used, membranes were blocked in TBST with 2% BSA, incubated with HRP-conjugated secondary antibodies, washed, and developed with ECL developer (Cytex) before imaging on the ChemiDoc.

### Imaging T cell migration on ICAM-1

Ibidi 8-well, glass-bottomed chamber slides (cat # 80807) were coated with ICAM-1 at 2 μg/mL overnight at 4°C. Activated CD4^+^ T cells were washed and resuspended in Leibovitz’s L-15 medium (Gibco) containing 2 mg/mL D-glucose and 0.1% FBS. ICAM-1-coated slides were washed with L-15 medium and incubated for 20 min at 37°C before 5×10^4^ cells were added to each well. Slides were incubated at 37°C for 20 min before being gently washed once to remove non-adherent cells and debris, then mounted for imaging. Imaging was performed on a Marianas system (3i) consisting of a Ziess Axio Observer 7 equipped with an X-Cite mini+ light source (Excelitas) and a Prime BSI Express CMOS camera (Photometrics) enclosed in an environmental chamber (Okolab). Images were collected every 30 sec for 10 min using 20ˣ objective at 37°C with Slidebook software (3i) and exported to ImageJ for analysis. Individual cells were tracked using the manual tracking plugin to generate xy coordinates for all motile cells. Cells were considered motile if they moved more than two full cell lengths during the entire imaging period. We also only included cells that stayed in the field of view for the entire imaging period. The xy coordinates were analyzed using the open-source software Migrate3D ^41^ to determine a variety of migratory parameters, including MSD, track length, Euclidean distance/displacement, speed, and directionality.

### Transendothelial migration assay

3B-11 endothelial cells were grown to confluency on μ-slide 8-well ibiTreat ibidi slides (cat #80806) in 400 μL of 3B-11 medium (see cell culture section) at 37°C. 24 h before imaging cells were treated with 50 ng/mL mouse TNFα or left untreated. Activated CD4^+^ T cells were washed and resuspended in warm Leibovitz’s L-15 medium (Gibco) containing 2 mg/mL D-glucose and 0.1% FBS. Confluent 3B-11 cells were washed once with the L-15 medium to remove debris and then mounted at 37°C for imaging. After 10 min of incubation at 37°C, 2×10^3^ T cells were added to the well. T cells were given 3 min to settle before imaging. Imaging was performed on a Marianas system (3i) consisting of a Ziess Axio Observer 7 equipped with an X-Cite mini+ light source (Excelitas) and a Prime BSI Express CMOS camera (Photometrics) enclosed in an environmental chamber (Okolab). Time-lapse images were collected every 30 s for 10 min using a 20x objective at 37°C with Slidebook software (3i) and exported to ImageJ for analysis. T cells were considered motile if they moved more than one full cell length during the imaging period. Migratory cells were scored based on whether they transmigrated through the endothelium or remained on the apical side.

### Statistical analysis

Statistics were calculated using GraphPad Prism 9 and Microsoft Excel. When only two groups were compared a two-tailed unpaired t-test was used. When more than two groups were compared, a one-way ANOVA was performed followed by post-hoc test to correct for multiple comparisons. \**p*<0.05; \*\**p*<0.01; \*\*\**p*<0.001; \*\*\*\**p*<0.0001.

### SUPPLEMENTARY FIGURE LEGENDS

**Movie 1-T cells migrating on untreated 3B-11 endothelial cells.**

Primary T cells were allowed to migrate on untreated 3B-11 endothelial cells and imaged every 30 seconds for 1 h total. The first 30 min is shown. The field is 270 μm x 270 μm.

**Movie 2-T cells migrating on TNFα 3B-11 endothelial cells**

Primary T cells were allowed to migrate on TNFα-treated 3B-11 endothelial cells and imaged every 30 seconds for 1 h total. The first 30 min is shown. The field is 270 μm x 270 μm.

## REFERENCES

1 Alcaide, P. Mechanisms Regulating T Cell-Endothelial Cell Interactions. Cold Spring Harb Perspect Med 12 (2022). 10.1101/cshperspect.a041170

2 Nourshargh, S. & Alon, R. Leukocyte migration into inflamed tissues. Immunity 41, 694–707 (2014). 10.1016/j.immuni.2014.10.008

3 Vestweber, D. How leukocytes cross the vascular endothelium. Nat Rev Immunol 15, 692–704 (2015). 10.1038/nri3908

4 Gronloh, M. L. B., van der Meer, W. J., Tebbens, M. E. & van Buul, J. D. Leukocyte transendothelial migration hotspots at a glance. J Cell Sci 138 (2025). 10.1242/jcs.263862

5 Park, E. J. et al. Distinct roles for LFA-1 affinity regulation during T-cell adhesion, diapedesis, and interstitial migration in lymph nodes. Blood 115, 1572–1581 (2010). 10.1182/blood-2009-08-237917

6 Gorina, R., Lyck, R., Vestweber, D. & Engelhardt, B. beta2 integrin-mediated crawling on endothelial ICAM-1 and ICAM-2 is a prerequisite for transcellular neutrophil diapedesis across the inflamed blood-brain barrier. J Immunol 192, 324–337 (2014). 10.4049/jimmunol.1300858

7 Porter, J. C. & Hall, A. Epithelial ICAM-1 and ICAM-2 regulate the egression of human T cells across the bronchial epithelium. FASEB J 23, 492–502 (2009). 10.1096/fj.08-115899

8 Shulman, Z. et al. Lymphocyte crawling and transendothelial migration require chemokine triggering of high-affinity LFA-1 integrin. Immunity 30, 384–396 (2009). 10.1016/j.immuni.2008.12.020

9 Alter, A. et al. Determinants of human B cell migration across brain endothelial cells. J Immunol 170, 4497–4505 (2003). 10.4049/jimmunol.170.9.4497

10 Rahman, A. & Fazal, F. Hug tightly and say goodbye: role of endothelial ICAM-1 in leukocyte transmigration. Antioxid Redox Signal 11, 823–839 (2009). 10.1089/ars.2008.2204

11 Kelleher, D., Murphy, A. & Cullen, D. Leukocyte function-associated antigen 1 (LFA-1) is a signaling molecule for cytoskeletal changes in a human T cell line. Eur J Immunol 20, 2351–2354 (1990). 10.1002/eji.1830201028

12 Porter, J. C., Bracke, M., Smith, A., Davies, D. & Hogg, N. Signaling through integrin LFA-1 leads to filamentous actin polymerization and remodeling, resulting in enhanced T cell adhesion. J Immunol 168, 6330–6335 (2002). 10.4049/jimmunol.168.12.6330

13 Smith, A., Bracke, M., Leitinger, B., Porter, J. C. & Hogg, N. LFA-1-induced T cell migration on ICAM-1 involves regulation of MLCK-mediated attachment and ROCK-dependent detachment. J Cell Sci 116, 3123–3133 (2003). 10.1242/jcs.00606

14 Anderson, N. R., Buffone, A., Jr. & Hammer, D. A. T lymphocytes migrate upstream after completing the leukocyte adhesion cascade. Cell Adh Migr 13, 163–168 (2019). 10.1080/19336918.2019.1587269

15 Hornung, A. et al. A Bistable Mechanism Mediated by Integrins Controls Mechanotaxis of Leukocytes. Biophys J 118, 565–577 (2020). 10.1016/j.bpj.2019.12.013

16 Anderson, D. C. et al. The severe and moderate phenotypes of heritable Mac-1, LFA-1 deficiency: their quantitative definition and relation to leukocyte dysfunction and clinical features. J Infect Dis 152, 668–689 (1985). 10.1093/infdis/152.4.668

17 Anderson, D. C. & Springer, T. A. Leukocyte adhesion deficiency: an inherited defect in the Mac-1, LFA-1, and p150,95 glycoproteins. Annu Rev Med 38, 175–194 (1987). 10.1146/annurev.me.38.020187.001135

18 Hogg, N. & Bates, P. A. Genetic analysis of integrin function in man: LAD-1 and other syndromes. Matrix Biol 19, 211–222 (2000). 10.1016/s0945-053x(00)00066-4

19 Cheung, S. M. & Ostergaard, H. L. Pyk2 Controls Integrin-Dependent CTL Migration through Regulation of De-Adhesion. J Immunol 197, 1945-1956 (2016). 10.4049/jimmunol.1501505

20 Roy, N. H., MacKay, J. L., Robertson, T. F., Hammer, D. A. & Burkhardt, J. K. Crk adaptor proteins mediate actin-dependent T cell migration and mechanosensing induced by the integrin LFA-1. Sci Signal 11 (2018). 10.1126/scisignal.aat3178

21 Ni, H. T., Deeths, M. J. & Mescher, M. F. LFA-1-mediated costimulation of CD8+ T cell proliferation requires phosphatidylinositol 3-kinase activity. J Immunol 166, 6523–6529 (2001). 10.4049/jimmunol.166.11.6523

22 Sanchez-Martin, L. et al. Signaling through the leukocyte integrin LFA-1 in T cells induces a transient activation of Rac-1 that is regulated by Vav and PI3K/Akt-1. J Biol Chem 279, 16194–16205 (2004). 10.1074/jbc.M400905200

23 Verma, N. K. et al. LFA-1/ICAM-1 Ligation in Human T Cells Promotes Th1 Polarization through a GSK3beta Signaling-Dependent Notch Pathway. J Immunol 197, 108-118 (2016). 10.4049/jimmunol.1501264

24 Cain, R. J. & Ridley, A. J. Phosphoinositide 3-kinases in cell migration. Biol Cell 101, 13–29 (2009). 10.1042/BC20080079

25 Saarikangas, J., Zhao, H. & Lappalainen, P. Regulation of the actin cytoskeleton-plasma membrane interplay by phosphoinositides. Physiol Rev 90, 259–289 (2010). 10.1152/physrev.00036.2009

26 Hou, X. et al. Phosphoinositide signaling at the cytoskeleton in the regulation of cell dynamics. Cell Death Dis 16, 296 (2025). 10.1038/s41419-025-07616-x

27 Katoh, K. Signal Transduction Mechanisms of Focal Adhesions: Src and FAK-Mediated Cell Response. Front Biosci (Landmark Ed) 29, 392 (2024). 10.31083/j.fbl2911392

28 Playford, M. P. & Schaller, M. D. The interplay between Src and integrins in normal and tumor biology. Oncogene 23, 7928–7946 (2004). 10.1038/sj.onc.1208080

29 Mitra, S. K. & Schlaepfer, D. D. Integrin-regulated FAK-Src signaling in normal and cancer cells. Curr Opin Cell Biol 18, 516–523 (2006). 10.1016/j.ceb.2006.08.011

30 Meng, F. & Lowell, C. A. A beta 1 integrin signaling pathway involving Src-family kinases, Cbl and PI-3 kinase is required for macrophage spreading and migration. EMBO J 17, 4391–4403 (1998). 10.1093/emboj/17.15.4391

31 He, Y. et al. The non-receptor tyrosine kinase Lyn controls neutrophil adhesion by recruiting the CrkL-C3G complex and activating Rap1 at the leading edge. J Cell Sci 124, 2153–2164 (2011). 10.1242/jcs.078535

32 Lowell, C. A. & Berton, G. Resistance to endotoxic shock and reduced neutrophil migration in mice deficient for the Src-family kinases Hck and Fgr. Proc Natl Acad Sci U S A 95, 7580–7584 (1998). 10.1073/pnas.95.13.7580

33 Zhang, H., Meng, F., Chu, C. L., Takai, T. & Lowell, C. A. The Src family kinases Hck and Fgr negatively regulate neutrophil and dendritic cell chemokine signaling via PIR-B. Immunity 22, 235–246 (2005). 10.1016/j.immuni.2005.01.004

34 Fumagalli, L., Zhang, H., Baruzzi, A., Lowell, C. A. & Berton, G. The Src family kinases Hck and Fgr regulate neutrophil responses to N-formyl-methionyl-leucyl-phenylalanine. J Immunol 178, 3874–3885 (2007). 10.4049/jimmunol.178.6.3874

35 Palacios, E. H. & Weiss, A. Function of the Src-family kinases, Lck and Fyn, in T-cell development and activation. Oncogene 23, 7990–8000 (2004). 10.1038/sj.onc.1208074

36 Molina, T. J. et al. Profound block in thymocyte development in mice lacking p56lck. Nature 357, 161–164 (1992). 10.1038/357161a0

37 Stein, P. L., Lee, H. M., Rich, S. & Soriano, P. pp59fyn mutant mice display differential signaling in thymocytes and peripheral T cells. Cell 70, 741–750 (1992). 10.1016/0092-8674(92)90308-y

38 van Oers, N. S., Lowin-Kropf, B., Finlay, D., Connolly, K. & Weiss, A. alpha beta T cell development is abolished in mice lacking both Lck and Fyn protein tyrosine kinases. Immunity 5, 429–436 (1996). 10.1016/s1074-7613(00)80499-9

39 Huang, B., Johansen, K. H. & Schwartzberg, P. L. Efficient CRISPR/Cas9-Mediated Mutagenesis in Primary Murine T Lymphocytes. Curr Protoc Immunol 124, e62 (2019). 10.1002/cpim.62

40 Schindelin, J. et al. Fiji: an open-source platform for biological-image analysis. Nat Methods 9, 676–682 (2012). 10.1038/nmeth.2019

41 Emily R. Mynar, M. W. K., Jonah Harris, Safwan Wshah, Markus Thali, Menelaos Symeonides. Migrate3D: Software for simplified post-tracking analysis of 2D and 3D object movement data. Research Square (2026). 10.21203/rs.3.rs-2451513/v3

42 Vadas, O., Burke, J. E., Zhang, X., Berndt, A. & Williams, R. L. Structural basis for activation and inhibition of class I phosphoinositide 3-kinases. Sci Signal 4, re2 (2011). 10.1126/scisignal.2002165

43 Lo, W. L. et al. Lck promotes Zap70-dependent LAT phosphorylation by bridging Zap70 to LAT. Nat Immunol 19, 733–741 (2018). 10.1038/s41590-018-0131-1

44 Kochl, R. et al. WNK1 kinase balances T cell adhesion versus migration in vivo. Nat Immunol 17, 1075–1083 (2016). 10.1038/ni.3495

45 Raab, M. et al. LFA-1 activates focal adhesion kinases FAK1/PYK2 to generate LAT-GRB2-SKAP1 complexes that terminate T-cell conjugate formation. Nat Commun 8, 16001 (2017). 10.1038/ncomms16001

46 Aasheim, H. C., Delabie, J. & Finne, E. F. Ephrin-A1 binding to CD4+ T lymphocytes stimulates migration and induces tyrosine phosphorylation of PYK2. Blood 105, 2869–2876 (2005). 10.1182/blood-2004-08-2981

47 Soede, R. D., Wijnands, Y. M., Van Kouteren-Cobzaru, I. & Roos, E. ZAP-70 tyrosine kinase is required for LFA-1-dependent T cell migration. J Cell Biol 142, 1371–1379 (1998). 10.1083/jcb.142.5.1371

48 Evans, R., Lellouch, A. C., Svensson, L., McDowall, A. & Hogg, N. The integrin LFA-1 signals through ZAP-70 to regulate expression of high-affinity LFA-1 on T lymphocytes. Blood 117, 3331–3342 (2011). 10.1182/blood-2010-06-289140

49 Haap-Hoff, A. et al. RNAi library screening reveals Gbeta1, Casein Kinase 2 and ICAP-1 as novel regulators of LFA-1-mediated T cell polarity and migration. Immunol Cell Biol 103, 73–92 (2025). 10.1111/imcb.12838

50 Bernardini, G., Kim, J. Y., Gismondi, A., Butcher, E. C. & Santoni, A. Chemoattractant induces LFA-1 associated PI 3K activity and cell migration that are dependent on Fyn signaling. FASEB J 19, 1305–1307 (2005). 10.1096/fj.04-3352fje

51 Inngjerdingen, M., Torgersen, K. M. & Maghazachi, A. A. Lck is required for stromal cell-derived factor 1 alpha (CXCL12)-induced lymphoid cell chemotaxis. Blood 99, 4318-4325 (2002). 10.1182/blood.v99.12.4318

52 Llavero, F., Artaso, A., Lacerda, H. M., Parada, L. A. & Zugaza, J. L. Lck/PLCgamma control migration and proliferation of interleukin (IL)-2-stimulated T cells via the Rac1 GTPase/glycogen phosphorylase pathway. Cell Signal 28, 1713–1724 (2016). 10.1016/j.cellsig.2016.07.014

53 Okabe, S., Fukuda, S. & Broxmeyer, H. E. Src kinase, but not the src kinase family member p56lck, mediates stromal cell-derived factor 1alpha/CXCL12-induced chemotaxis of a T cell line. J Hematother Stem Cell Res 11, 923–928 (2002). 10.1089/152581602321080583

54 Ryan, T. C., Cruikshank, W. W., Kornfeld, H., Collins, T. L. & Center, D. M. The CD4-associated tyrosine kinase p56lck is required for lymphocyte chemoattractant factor-induced T lymphocyte migration. J Biol Chem 270, 17081–17086 (1995). 10.1074/jbc.270.29.17081

55 Zaman, S. N., Resek, M. E. & Robbins, S. M. Dual acylation and lipid raft association of Src-family protein tyrosine kinases are required for SDF-1/CXCL12-mediated chemotaxis in the Jurkat human T cell lymphoma cell line. J Leukoc Biol 84, 1082–1091 (2008). 10.1189/jlb.1007698

56 Phan, T. X., Jaruga, B., Pingle, S. C., Bandyopadhyay, B. C. & Ahern, G. P. Intrinsic Photosensitivity Enhances Motility of T Lymphocytes. Sci Rep 6, 39479 (2016). 10.1038/srep39479

57 Caillier, A., Oleksyn, D., Fowell, D. J., Miller, J. & Oakes, P. W. T cells use focal adhesions to pull themselves through confined environments. J Cell Biol 223 (2024). 10.1083/jcb.202310067

58 Sarantos, M. R. et al. Transmigration of neutrophils across inflamed endothelium is signaled through LFA-1 and Src family kinase. J Immunol 181, 8660–8669 (2008). 10.4049/jimmunol.181.12.8660

59 Medina, I. et al. Hck/Fgr Kinase Deficiency Reduces Plaque Growth and Stability by Blunting Monocyte Recruitment and Intraplaque Motility. Circulation 132, 490–501 (2015). 10.1161/CIRCULATIONAHA.114.012316

60 Shulman, Z. et al. Transendothelial migration of lymphocytes mediated by intraendothelial vesicle stores rather than by extracellular chemokine depots. Nat Immunol 13, 67–76 (2011). 10.1038/ni.2173

61 Carman, C. V. et al. Transcellular diapedesis is initiated by invasive podosomes. Immunity 26, 784–797 (2007). 10.1016/j.immuni.2007.04.015

62 Schaeuble, K., Hauser, M. A., Singer, E., Groettrup, M. & Legler, D. F. Cross-talk between TCR and CCR7 signaling sets a temporal threshold for enhanced T lymphocyte migration. J Immunol 187, 5645–5652 (2011). 10.4049/jimmunol.1101850

63 Taichman, R., Merida, I., Torigoe, T., Gaulton, G. N. & Reed, J. C. Evidence that protein tyrosine kinase p56-Lck regulates the activity of phosphatidylinositol-3’-kinase in interleukin-2-dependent T-cells. J Biol Chem 268, 20031–20036 (1993).

64 Carrera, A. C., Rodriguez-Borlado, L., Martinez-Alonso, C. & Merida, I. T cell receptor-associated alpha-phosphatidylinositol 3-kinase becomes activated by T cell receptor cross-linking and requires pp56lck. J Biol Chem 269, 19435–19440 (1994).

65 von Willebrand, M., Baier, G., Couture, C., Burn, P. & Mustelin, T. Activation of phosphatidylinositol-3-kinase in Jurkat T cells depends on the presence of the p56lck tyrosine kinase. Eur J Immunol 24, 234–238 (1994). 10.1002/eji.1830240137

66 Raab, M. et al. p56Lck and p59Fyn regulate CD28 binding to phosphatidylinositol 3-kinase, growth factor receptor-bound protein GRB-2, and T cell-specific protein-tyrosine kinase ITK: implications for T-cell costimulation. Proc Natl Acad Sci U S A 92, 8891–8895 (1995). 10.1073/pnas.92.19.8891

67 de Souza, A. J. et al. T cell Ig and mucin domain-1-mediated T cell activation requires recruitment and activation of phosphoinositide 3-kinase. J Immunol 180, 6518–6526 (2008). 10.4049/jimmunol.180.10.6518

68 Roy, N. H. et al. LFA-1 signals to promote actin polymerization and upstream migration in T cells. J Cell Sci 133 (2020). 10.1242/jcs.248328

